# TIGIT expression dictates the immunosuppressive reprogramming of myeloid cells in glioblastoma

**DOI:** 10.1101/2025.04.24.648993

**Authors:** Mohammad Asad, Julio Inocencio, Stefan Mitrasinovic, Minori Aoki, Celina Crisman, Patrick Lasala, Emad Eskandar, Chandan Guha, XingXing Zang, Ian F. Parney, Benjamin T. Himes

## Abstract

Glioblastoma (GBM) is a deadly brain cancer with near-universal recurrence despite maximal treatment for which new innovations are sorely needed. Immunotherapy has yet to make significant gains in GBM treatment despite revolutionizing other cancer therapies, due in part to GBM-mediated immune suppression. This immune derangement proceeds through several mechanisms, but increasing evidence points to critical roles for tumor-derived extracellular vesicles (EVs) and immunosuppressive myeloid cells as key factors in this process. In the present study, we demonstrate broad expression of TIGIT across myeloid cell populations in the GBM microenvironment, a finding recapitulated by conditioning healthy monocytes with GBM-derived EVs. Further, knockdown of TIGIT expression reduced the immunosuppressive polarization of monocytes, resulting in improvement in T cell function. This finding proceeded in an NLRP3-dependent manner, with substantial co-localization of TIGIT and NLRP3 expression prior to knockdown. These findings point to a novel role for TIGIT expression in diverse myeloid cells in the GBM microenvironment as a marker of immunosuppressive activity and further indicate a hierarchy of immunomodulatory protein activity in these myeloid cells, with TIGIT knockdown unmasking the pro-inflammatory activity of NLRP3. This study bolsters understanding of the immunosuppressive complexities of myeloid cells in the GBM microenvironment, while lending further support to prevention or attenuation of immunosuppressive myeloid cell activity as a means of restoring immune function in GBM.

**Graphical abstract:** (Created in BioRender. Asad, M. (2025) https://BioRender.com/euiljoq

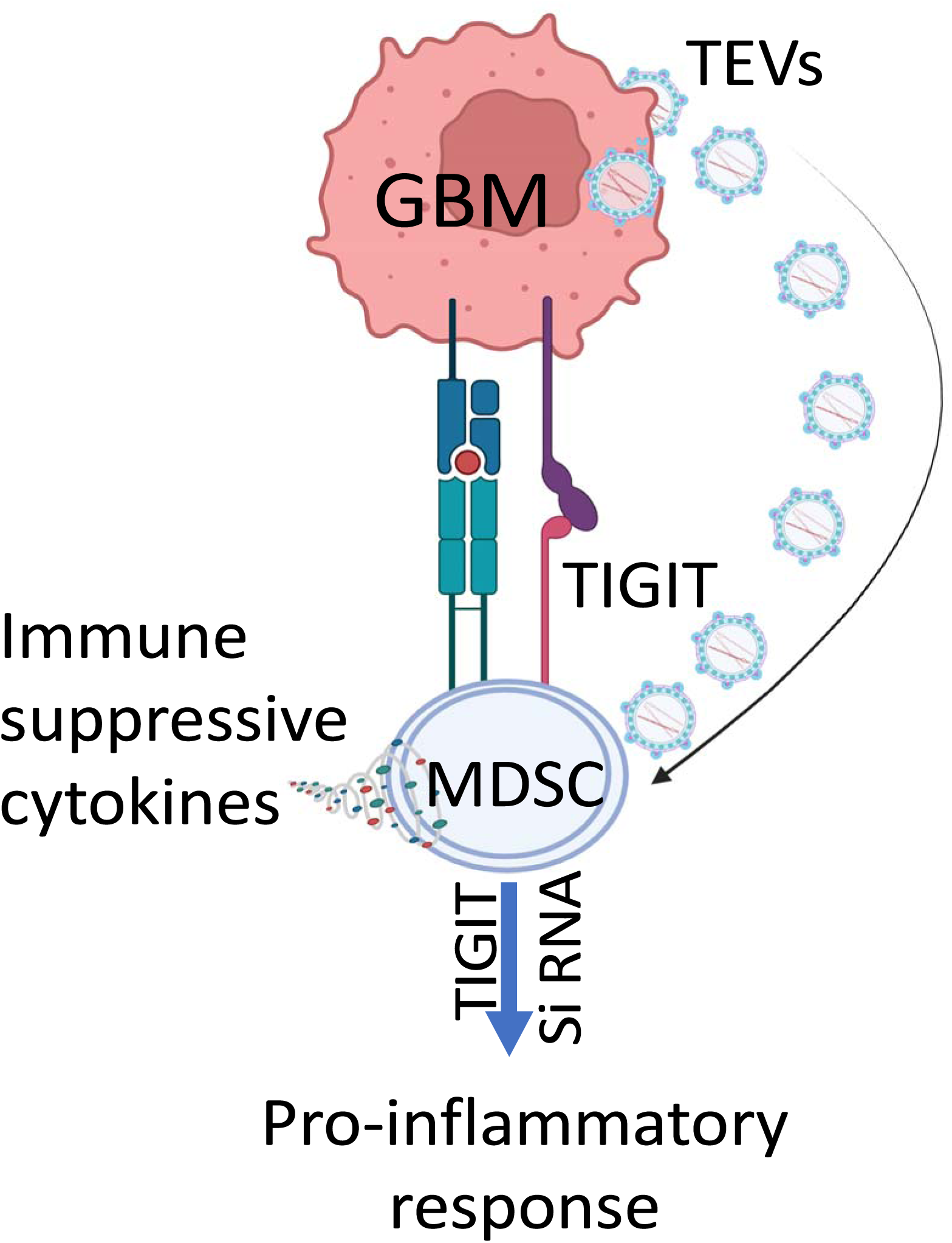

**Key Points:**

1. Tumor-mediated immune suppression is a key barrier to the development of effective immunotherapies for GBM.
2. TIGIT is broadly expressed in myeloid cells within the GBM microenvironment and can be induced by GBM-derived extracellular vesicles.
3. Knockdown of TIGIT reduces immunosuppressive polarization of monocytes and enhances T cell activity via NLRP3 signaling, implicating TIGIT expression as a targetable modifier of immunosuppressive activity in GBM-associated myeloid cells.

**Importance of the Study:** Glioblastoma (GBM) remains a formidable clinical challenge, with poor prognosis and limited response to current immunotherapies. This study uncovers a novel immunosuppressive axis involving TIGIT expression in myeloid cells, which are key players in the GBM tumor microenvironment. By demonstrating GBM-derived extracellular vesicles induce TIGIT in healthy monocytes and TIGIT knockdown diminishes immunosuppressive polarization in an NLRP3-dependent manner, this study highlights TIGIT as both a marker and modulator of immune dysfunction in GBM. These findings introduce a functional hierarchy of immunoregulatory proteins in tumor-associated myeloid cells, positioning TIGIT as a potential checkpoint target. Restoring immune activity by disrupting this axis could enhance the efficacy of immunotherapy in GBM. Thus, this research not only advances our understanding of tumor-induced immune suppression but also opens a promising therapeutic avenue to reinvigorate anti-tumor immunity in a cancer type historically resistant to immunotherapeutic approaches.

## Introduction

Glioblastoma (GBM) is the most common and deadly intrinsic brain tumor in adults. Current standard of care consists of maximal safe surgical resection followed by radiation and temozolomide chemotherapy, in a protocol essentially unchanged for two decades (*1*). Some recent innovations, such as tumor treating fields, have shown modest promise but have not led to increased median overall survival beyond 15 months in population-based studies (*2*). Immunotherapy, including immune checkpoint blockade, has been transformative in cancer care in recent years, but has failed to improve outcomes in GBM (*3–6*). The profound immunosuppression found in GBM patients plays a critical role in these failures, but its underlying mechanisms remain poorly understood (*6, 7*).

Increasing evidence points to a crucial role for myeloid cells, which can comprise the bulk of the tumor immune infiltrate, as mediators of immune suppression, preventing the antitumor effects of T cells (*7, 8*). These myeloid cells are heterogenous with and include a number of distinct populations, including myeloid-derived suppressor cells (MDSCs) and non-classical monocytes (NCM) (*8–12*). These diverse populations are united by the common trait of facilitating immune suppression, whether by release of immunosuppressive cytokines or expression of surface ligands that facilitate exhaustion of effector cells, including T cells (*7, 13*).

The mechanisms by which GBM induces immunosuppression are multiple and an active field of study, but an increasing body of work points to a central role of tumor-derived extracellular vesicles (EVs) (*7, 10, 14, 15*). These complex, multimodal signaling molecules, are shed by all cells and play a critical role in cancer signaling, particularly in immune evasion (*16*). In GBM, work by our group and others has pointed to a potentially pivotal role of EVs in tumor-mediated immune suppression, including the induction of immunosuppressive myeloid cells (*10, 14, 17*). Thus, the means by which GBM induces the immunosuppressive polarization of myeloid cells represents a potentially potent therapeutic axis for mitigation of GBM-mediated immune suppression.

In the present study, we examine the immune phenotype of myeloid cells in the GBM tumor microenvironment at the time of surgical resection using ultrasonic aspirate processed directly from the operating room to identify immunomodulatory markers expressed across diverse myeloid cell populations that may represent therapeutically significant targets across diverse cell populations. Interestingly, we found high rates of TIGIT (T cell immunoreceptor with immunoglobulin and ITIM domain). TIGIT is increasingly recognized as playing a critical role in limiting adaptive and innate immunity and is canonically understood as a marker for T cell exhaustion. Recent data demonstrates TIGIT expression across an array of myeloid cells in the *in-situ* GBM microenvironment (*18–20*). These findings were recapitulated using our *in vitro* system of immunosuppressive monocyte induction, whereupon healthy monocytes are subjected to GBM-EVs collected from patient-derived cell lines. Upon knockdown of TIGIT expression in monocytes, we found cells underwent a more pro-inflammatory immune polarization in response to EV conditioning. TIGIT knockdown unmasked high levels of co-expressed NLRP3 and increased levels of IL-13 expression. This resulted in a rescue of T cell proliferation upon subsequent co-culture, indicating a level of functional restoration of immune function. These results illustrate the potentially crucial role of an immunomodulatory hierarchy in myeloid cells in the tumor microenvironment, the understanding of which may prove critical for reversal of tumor-mediated immune suppression.

## Results

### Immunophenotyping of myeloid cells in the GBM microenvironment reveals broad TIGIT expression

GBM is an intrinsic infiltrative tumor, with the radiographic tumor bulk comprised of heterogenous tumor cells, immune infiltrate, and normal brain cells. Surgical resection of GBM entails removing as much of the lesion as safely possible with a range of surgical techniques, frequently including liberal use of an ultrasonic aspirator, which breaks up the bulk tumor and allows for resection without excess trauma due to heat or retraction on the surrounding brain. The ultrasonic aspirate from this approach thus consists of all of the constituent cells of the bulk tumor and is thus a valuable source of material for study of the *in situ* tumor microenvironment, as well as tumor cells for primary cell culture and establishment of patient-derived cell lines. Ultrasonic aspirate from 13 GBM patients was collected and the immune cell infiltrate profiled via high dimensional flow cytometry (Cytek Aurora, full panel and gating strategy available in **Table S1 and Fig. 1**). Analysis of a number of key immunomodulatory proteins were assessed across several known immunosuppressive monocyte populations, including monocytic myeloid-derived suppressor cells (mMDSCs, CD11b^+^CD15^-^CD14^+^HLA-DR^lo/-^) and non-classical monocytes (NCM, CD11b^+^CD15^-^ CD16^+^HLA-DR^+^PD-1^+^), that have been previously described as major populations of immunosuppressive myeloid cells in the GBM microenvironment (**Fig. 1A**) (*10, 14, 21*). Among the infiltrating immune cells, T cells constituted the largest population, followed by immunosuppressive monocytes and B cells. Subsequent analysis of immunomodulatory proteins in mMDSC and NCM populations revealed enriched expression of a few key immune modulatory proteins, such as VISTA, PD-L1 and B7-H3 (**Fig. 1B, C**). Interestingly, we found that TIGIT, typically thought of as a T cell exhaustion marker, was significantly elevated in elevated in immunosuppressive myeloid cell populations relative to other immunomodulatory markers. Despite the significant presence of PD-L1 (29.75±5.83%) and B7-H3 (50.34±9.093%) expressing cells in NCM, TIGIT^+^ cells predominated among the myeloid cell populations within the various suppressive monocytic subsets (64.27±7.99% and 39.8±4.366% in mMDSCs and NCM respectively). Analysis of The Cancer Genome Atlas (TCGA) data demonstrated enrichment of TIGIT expression across several GBM samples. Further, TCGA and GTEx datasets of human GBM was analyzed using Gene Expression Profiling Interactive Analysis. The TIGIT high group (top 20%, N= 128) indicated in red showed a significant worse overall survival than the TIGIT low group (the rest 80%, N= 130) in blue, p = 0.036 (**Fig. 1 D, E**). Thus, it appears that elevated TIGIT mRNA expression is associated with poor survival in human GBM patients.

**Figure 1:**
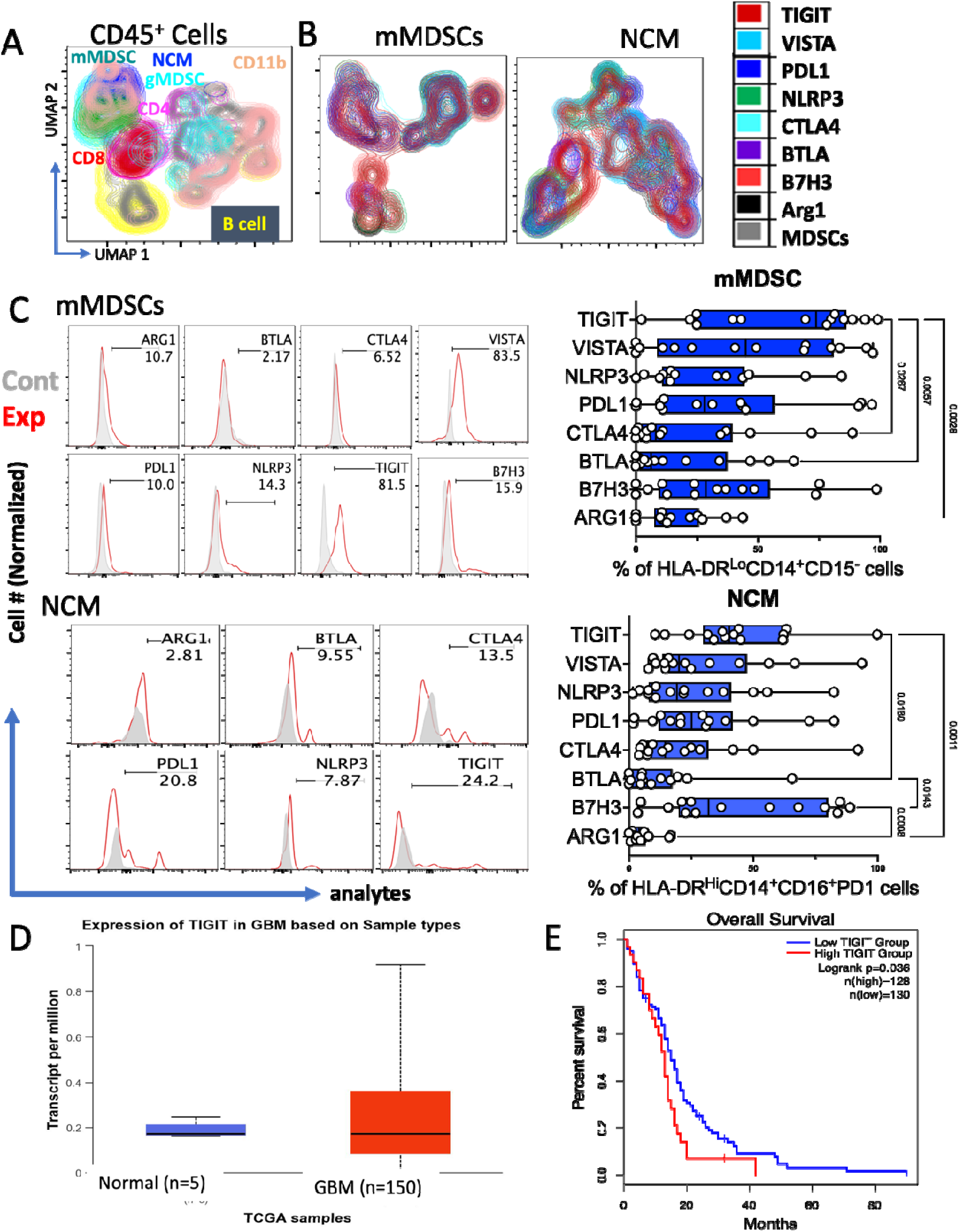
Expression of immune modulatory molecules by suppressive monocytes of GBM patients. For surveying the tumor-immune microenvironment in samples of ultrasonic aspirates collected during surgery, samples were processed for flow cytometric studies and the single cell suspensions, stained with various flow antibodies were run on Cytek aurora spectral flow cytometer. (**A**) UMAP analysis of CD45+ cells from GBM patients show various infiltrating immune cells. (**B**) Showing the cellular expression of immune modulatory molecules (TIGIT, PDL1, NLRP3, BTLA, CTLA4, ARG1) by infiltrating immune suppressive monocyte subsets, monocytic Myeloid Derived Suppressor Cells, mMDSCs (CD45^+^CD11b^+^CD15^-^HLA-DR^Lo^CD14 cells) and nonclassical monocytes (NCM, CD45^+^CD11b^+^CD15^-^HLA-DR^+^CD14^-^PD1^+^CD16 cells). (**C**) Histograms and a corresponding summary bar graph depicting the immune modulatory molecules expressing cell population in mMDSCs and NCM. (**D**) Box plot obtained from TCGA database illustrates the expression of TIGIT in normal and primary tumor samples obtained from GBM patients. (**E**) Kaplan-Meier curve of TCGA and GTEx datasets of human GBM, demonstrating the effect of TIGIT expression on patient survival in GBM. Data are presented as mean□±□s.e.m. and were analyzed by one-way analysis of variance (ANOVA) with Tukey’s multiple comparisons.

### GBM-EVs recapitulate myeloid cell immunosuppressive polarization *in vitro*

We next sought to validate these findings outside of the highly heterogenous context of the tumor microenvironment. Work by our group and others has pointed to a major role for GBM-derived extracellular vesicles (EVs) in the formation of immunosuppressive monocytes (*10*). We previously demonstrated that EVs collected from patient-derived GBM cell cultures can induce the formation of immunosuppressive monocytes, including mMDSCs and NCMs, when used to condition monocytes collected from healthy human blood donors. Using these techniques, we collected GBM-EVs from three patient-derived GBM cell lines (dBT114, dBT116, and dBT120). In order to collect EVs, GBM cells were cultured in serum-free conditions for 72hr and subjected to staged centrifugation as described above. Presence and sizing of EVs was confirmed by use of the NanoSight Nanotracker system (ZetaView, Particle Matrix, Germany) as well as the presence of the EV markers CD9 and CD63 via western blot (**Fig. 2A-C**). EV quantification was performed by BCA assay to quantify total protein. To induce the formation of immunosuppressive monocytes, 20ug of EVs was added to monocytes isolated from anonymous healthy blood donors (obtained from the New York Blood Bank). Monocytes were conditioned with EVs for 72h in hypoxic (1% O2) conditions prior to collection for flow cytometry analysis (panel included in **S2**). Consistent with our previously published results, EVs derived from dBT114, dBT116, and dBT120 all significantly induced the formation of both mMDSCs and NCM in this model system (**Fig. 2D**), which produce anti-inflammatory cytokines including IL-4 (**Fig. S2**).

**Figure 2:**
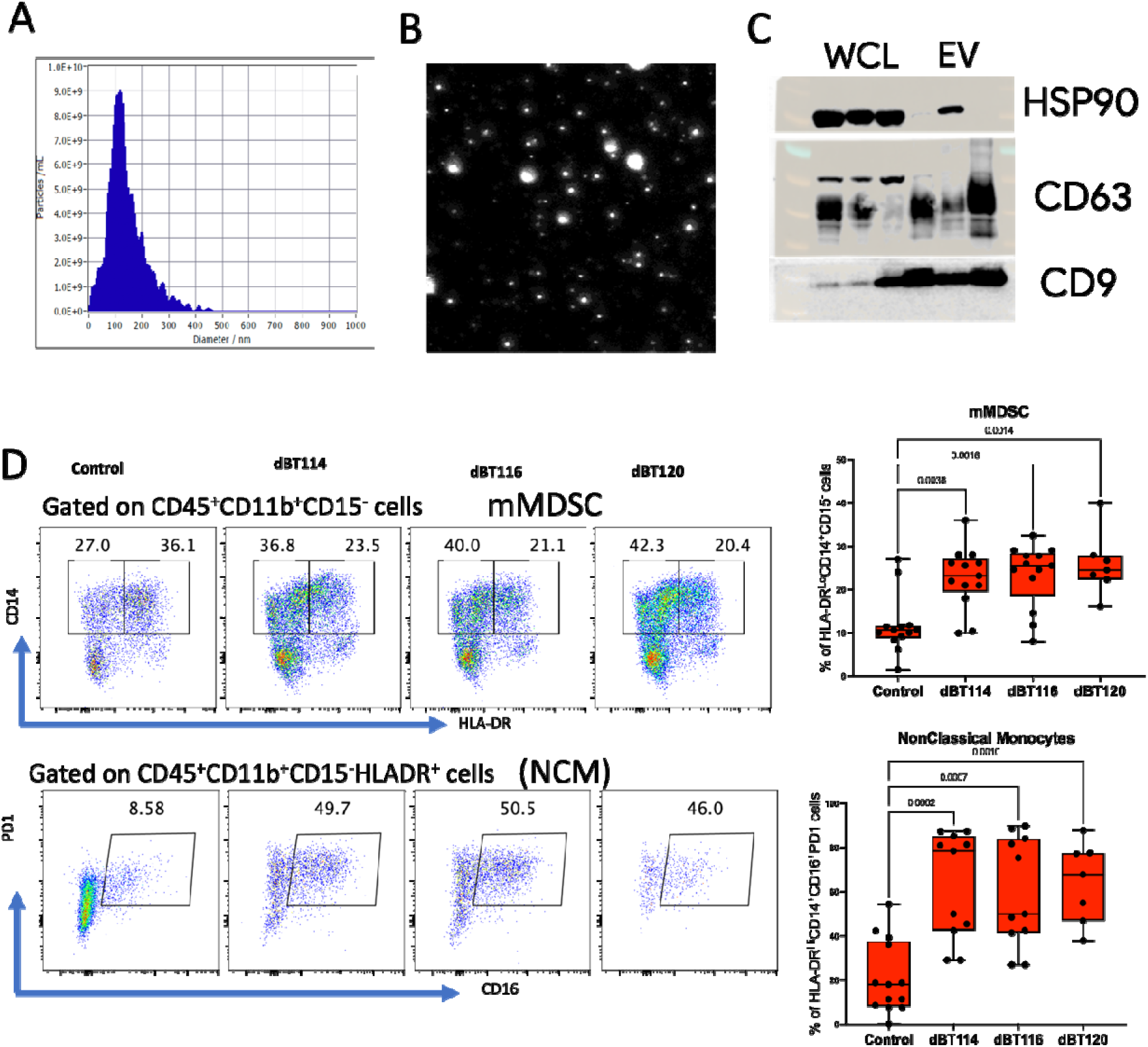
Isolation and characterization of tumor-derived extracellular vesicles (TEVs) and induction of immune suppressive monocytes. Extra cellular vesicles from human GBM primary cells lines were obtained by ultracentrifugation and characterized by Nano-sight and western blots. (**A**) Size distribution analysis of tumor-derived extracellular vesicles (TEVs) isolated from conditioned medium post ultracentrifugation. (**B**) Representative image depicting the shape and relative size of the heterogeneous TEVs, obtained from Nano-sight. (**C**) Western blot analysis of TEVs and whole cell lysate (WCL) for the presence of CD63, and CD9 as TEVs markers. (**D**) CD11b^+^ cells, isolated from healthy PBMCs and induced with control or different TEVs isolated from primary GBM cell lines (dBT114, dBT116 and dBT120) under hypoxic condition in serum free medium. After 72 hours, cells were collected and processed for flow cytometry and stained with various fluorochorm tagged antibodies. Flow cytometric analysis of monocytic Myeloid Derived Suppressor Cells (mMDSCs, CD45^+^CD11b^+^CD15^-^HLA-DR^Lo^CD14 cells and nonclassical monocytes (NCM, CD45^+^CD11b^+^CD15^-^HLA-DR^+^CD14^-^PD1^+^CD16 cells) isolated from healthy PBMCs and induced with control or different TEVs isolated from primary GBM cell lines. Data are presented as mean□±□s.e.m. and were analyzed by one-way analysis of variance (ANOVA) with Tukey’s multiple comparisons.

Using this model, we next employed a high-dimensional flow cytometry approach as employed for the ultrasonic aspirate samples to define the expression pattern of immunomodulatory proteins in this model. We found substantial similarities in immunomodulatory protein expression following EV-conditioning of monocytes, including consistent increases in TIGIT and NLRP3 expression (**Fig. 3A, B**). TIGIT expression was increased in both mMDSCs (73.53±6.27%, 68.33±8.66% and 63.99±8.11% respectively for the mentioned cell lines EVs, compared to unstimulated control, 4.68±2.83%) and NCM (85.17±3.91%, 79.26±9.17% and 60.68±10.52% respectively, compared to control, 11.3±3.3%) following treatment with EVs derived from all three GBM cell lines. We confirmed that TIGT expression in this model was present in monocytes and not from co-localized EVs via western blot demonstrating TIGIT expression in GBM whole cell lysate (WCL), but not in EVs (**Fig. 3C**). Further, immunofluorescent staining following EV-monocyte co-culture for 72h demonstrated robust TIGIT expression in monocytes conditioned with EVs, but not in untreated controls, demonstrating that hypoxic culture conditions, which have previously been shown to aid in MDSC induction, were insufficient to induce TIGIT expression (**Fig. 3D**).

**Figure 3:**
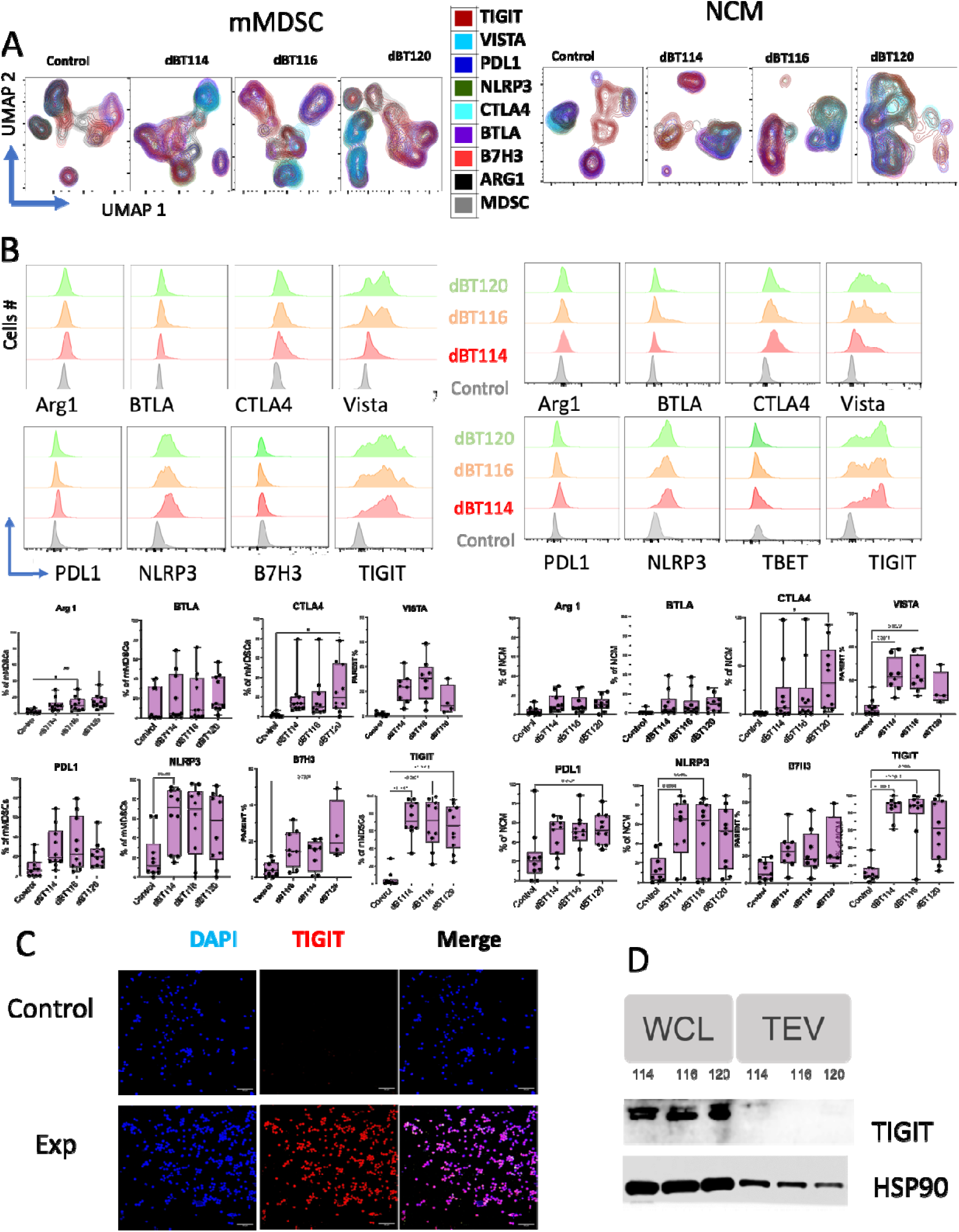
Expression of immune modulatory molecules on TEVs induced peripheral monocytic Myeloid Derived Suppressor Cells (mMDSC) & non-classical monocytes (NCM) CD11b^+^ cells, isolated from healthy peripheral blood and under hypoxic condition in a serum free medium, co-cultured for 72 hours with TEVs obtained from mentioned GBM cell lines. (**A**) TEVs induced CD11b^+^ monocytes were studied for the expression of various immune modulatory markers, TIGIT, PDL1, NLRP3, BTLA, CTLA4 & ARG1 on suppressive monocyte subsets (mMDSCs and NCMs). (**B**) Histograms depicting the expression levels of immune modulatory molecules on mMDSC and NCM populations cultured with control or different TEVs (dBT114, dBT116 and dBT120). The percentage of cells expressing each molecule within each subset is also shown correspondingly as a chart. (**C**) Immunofluorescence staining of TIGIT in suppressive monocytes treated with control or TEVs. (**D**) Western blot analysis of GBM whole cell lysate (WCL) for the presence of TIGIT. Data are presented as mean□±□s.e.m. and were analyzed by one-way analysis of variance (ANOVA) with Tukey’s multiple comparisons.

### TIGIT and key immunomodulatory proteins are co-expressed in EV-induced immunosuppressive monocyte populations

While there is some recent work describing TIGIT expression in myeloid cells (*20*), typically TIGIT has been viewed as a T cell exhaustion marker, and so it is not necessarily intuitive that high levels of TIGIT expression on immunomodulatory myeloid cells corresponds to downstream immunosuppressive functionality in myeloid cells. We, therefore next interrogated the immunomodulatory proteins and cytokines co-expressed on TIGIT+ immunosuppressive monocytes. Several other immune-checkpoint markers, including PD-L1 and VISTA, demonstrated high levels of co-expression on TIGIT positive mMDSCs and NCM following GBM-EV treatment (**Fig. 4A, B**, representative dot plots for co-localization included in **Fig. S3**). Interestingly, NLRP3, typically understood as a strong marker of pro-inflammatory activity important in the inflammasome pathway, demonstrated the strongest amount of co-expression. It is noteworthy that among all CD45^+^ cells, TIGIT^+^ cells showed a significantly higher expression of NLRP3^+^ cells (86.64±5.52%) compared to other immunomodulatory molecules, including Arg1 (0.45±0.14%), BTLA (2.2±0.94%), CTLA-4 (27.34±11.12%), and PD-L1 (48.09±4.41%) (Fig. S3c). This elevated NLRP3 expression was accompanied by a significant increase in IL-1β-producing cells (55.88±5%), an inflammatory cytokine associated with NLRP3, as well as IL-13-producing cells (98.78±.3%). These levels were notably higher than those of the proinflammatory cytokines IFNγ (7.3±2.65%) and IL-17 (1.63±0.63%), and the immunosuppressive cytokines IL-4 (5.9±1.3%), IL-5 (16.98±5%), and IL-10 (12.49±2.35%) (**Fig. S3**). In interrogating a panel of key cytokines, we found that the TH2 cytokine, IL-13 was co-expressed with TIGIT expression on immunosuppressive monocytes (**Fig. 4C, D**). Taken together, these findings suggest that an immunosuppressive phenotype prevails in the presence of co-expressed pro-inflammatory markers in EV-induced immunosuppressive monocytes, suggestive of an immunomodulatory protein hierarchy that may be leveraged to repolarize towards an anti-tumor immune response.

**Figure 4:**
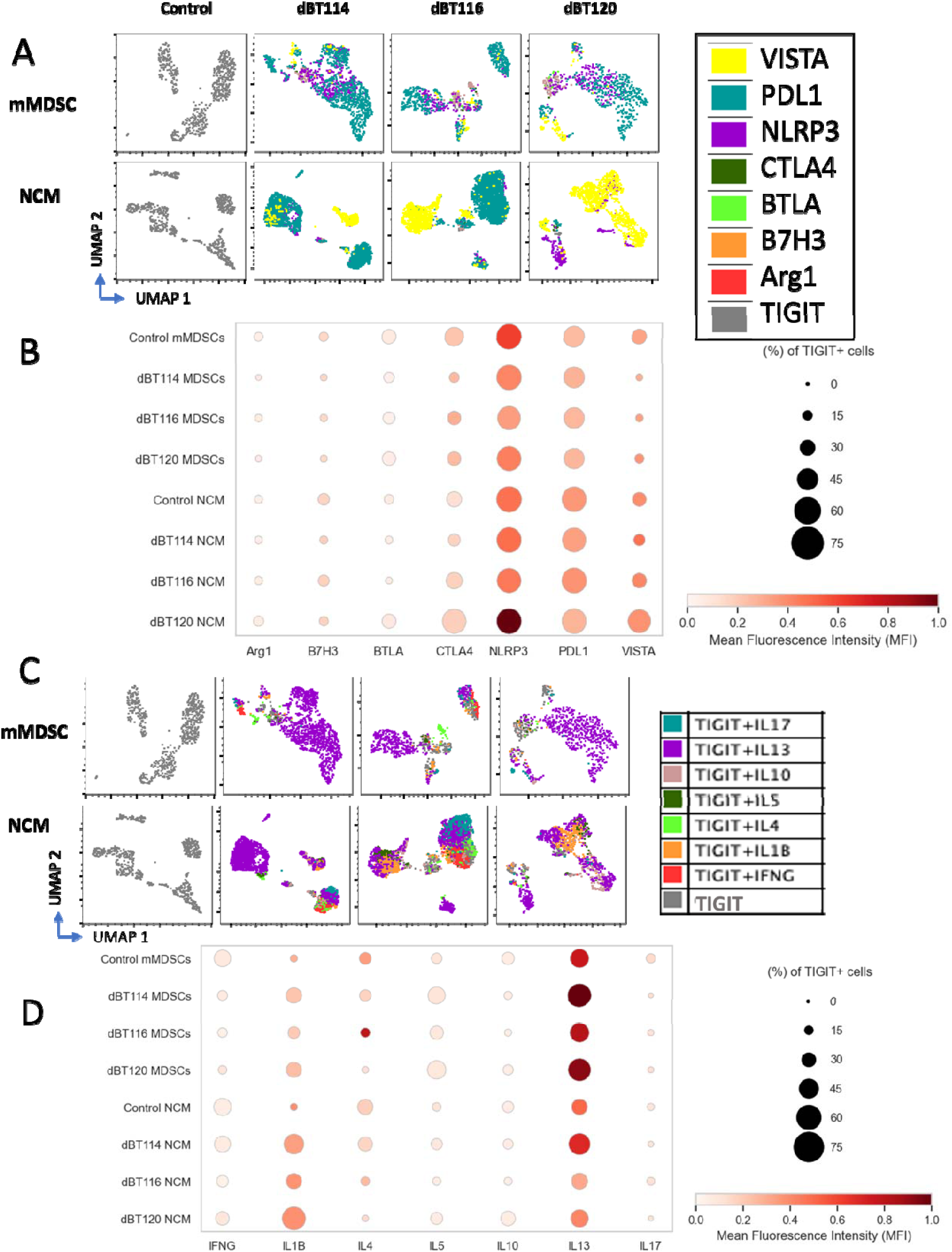
Immune response by TIGIT^+^ suppressive monocytes. Isolated peripheral blood CD11b^+^ monocytes were induced with TEVs in hypoxia chamber in serum free medium. After 72 hours of induction, cells were processed for flow cytometric studies. Single viable CD45^+^TIGIT cells were selected for the analysis. (**A**) UMAP analysis of mMDSCs and NCM treated with control or different TEV preparations (dBT114, dBT116, dBT120), highlighting the co-expression of various immune modulatory molecules (VISTA, PDL1, NLRP3, CTLA4, BTLA and ARG1) within TIGIT^+^ cell population. (**B**) Dotplot depicting the percentage of cells as a factor of size of the dot expressing different immune modulatory molecules within the TIGIT pockets of mMDSCs and NCMs treated with control or different EV preparations. Mean fluorescence intensity (MFI) shown corresponding to the color intensity of the dots for each analyte. (**C**) UMAP analysis of mMDSCs and NCM treated with control or different TEVs, highlighting the expression of various cytokines (IL17, IL13, IL10, IL5, IL4, IL18, IFNγ). (**D**) Dotplot depicting the percentage and MFI of TIGIT^+^ cells co-expressing different cytokines in mMDSCs and NCM treated with control or different EV preparations.

### TIGIT knockdown in EV-conditioned monocytes results in a pro-inflammatory phenotype

To test if the TIGIT driven immunomodulatory hierarchy regulates the NLRP3-mediated protective immune response against GBM, we next sought to determine if prevention of TIGIT expression on monocytes was sufficient to ‘unmask’ a pro-inflammatory phenotype in EV-induced immunosuppressive monocytes. Prior to EV conditioning, monocytes isolated from PBMCs were treated with TIGIT-directed siRNA to knock down TIGT expression. GBM-EVs were subsequently added for 72h of hypoxic co-culture as described above. TIGIT knockdown demonstrated immunomodulatory reprogramming in both mMDSC and NCM immunosuppressive monocyte populations (**Fig. 5**). While TIGIT knockdown in mMDSCs resulted in reduced expression of other key immunomodulatory markers including B7-H3 (11.6±3.39), PD-L1 (11.04±6.59), VISTA (12.3±2.05), and CTLA-4 (2.19±1.18, **Fig. 5B**) relative to scramble treated controls (32.73±6, 20.52±9.4, 29.46±5.6 and 18.12±8.1 respectively, **Fig. 5A**), this effect in NCMs did not reach a significant difference in any of the above mentioned immune regulatory proteins (B7-H3 15.72±8.09, PD-L1 16.49±4.37, VISTA 34.82±9.7 and CTLA-4 12.38±3.12, respectively) compared to scramble controls (41.55±5.19,37.04±10.35, 39.20±6.7 and 33.66±8.35, **Fig. 5C, D, Fig. S4**). Meanwhile, NLRP3 expressing cells remained significantly elevated following TIGIT knockdown in both populations (72.58±5.78 and 48.75±8.19 respectively compared to respective scrambles, 31.2±7.65 and 25.61 p<0.05, **Fig. 5B, D**).

**Figure 5.**
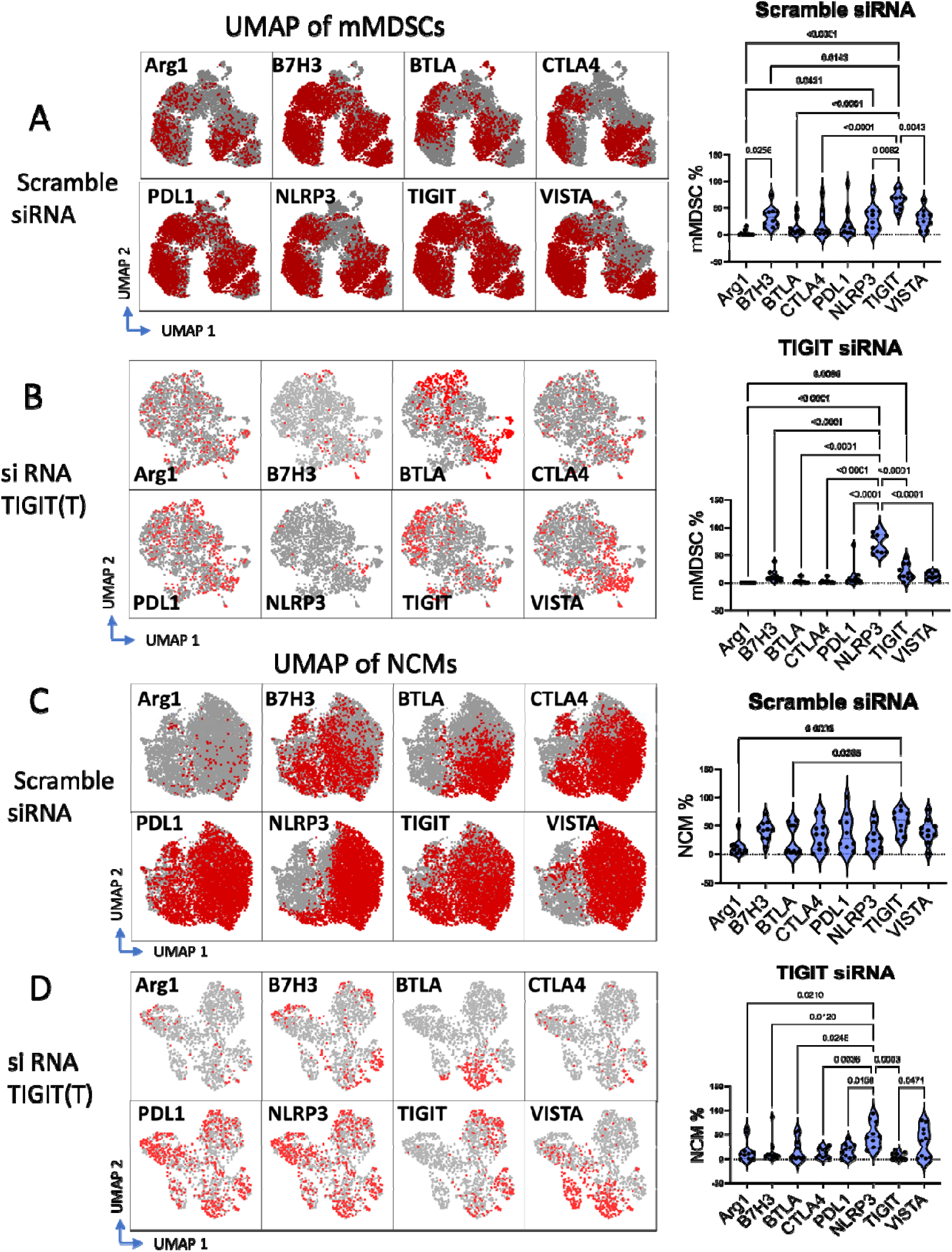
Immune modulatory molecules expressing Immune suppressive monocytes post TIGIT knockdown. Isolated CD11b^+^ Monocytic cells from healthy peripheral blood were knockdown by scramble or NLRP3-specific or TIGIT-specific siRNA and processed for flow cytometry post TEVs treatment for 72 hours under hypoxia in serum free condition. (**A and B**) Uniform Manifold Approximation and Projection (UMAP) plots showing the expression patterns of immune checkpoint molecules (Arg1, B7H3, BTLA, CTLA4, PD-L1, NLRP3, TIGIT, and VISTA) across different clusters by mMDSC population post TIGIT specific siRNA-(b) or scramble siRNA-(a) treated cells. Red color represents high expression, whereas gray indicates the total cellular population of mMDSCs. Violin plots depicting the percentage of mMDSCs expressing various immune modulatory molecules in TIGIT siRNA-treated and scramble siRNA-treated peripheral monocytes. (**C and D**) UMAP showing the expression of the same molecules by non-classical monocytes (NCM), after TIGIT siRNA-treated (d) and scramble siRNA-treated (c) peripheral monocytes. Violin plots depicting the percentage of NCMs expressing various immune modulatory molecules in TIGIT siRNA-treated and scramble siRNA-treated cells. Data are presented as mean□±□s.e.m. and were analyzed by one-way analysis of variance (ANOVA) with Tukey’s multiple comparisons.

We next examined the effect of TIGIT knockdown on cytokine expression in these immunosuppressive monocyte populations. This revealed a marked increase in IL17-expressing cells relative to scramble controls in both mMDSC and NCM populations (**Fig. 6B, D**). However, this enhanced IL17 expression was significantly higher in mMDSCs. Interestingly, increased IL17 expression was coupled with an attenuation of IL13 expression, in TIGIT downregulated monocyte subsets, which as mentioned above (**Fig. 4**) previously co-localized heavily with TIGIT^+^ immunosuppressive monocytes (**Fig. 6 A, C, Fig. S5**). Interestingly, IL13 expressing cells were observed to co-expressed with NLRP3^+^ cells within the mMDSCs and NCMs (**Fig. 5** and **6**). Furthermore, IL1β expressing cells also exhibited co-expression with NLRP3 in the NCM (**Fig. 5D** and **6D**). Collectively, these data indicate that TIGIT-dependent IL13 may play a role in regulating NLRP3-mediated inflammation.

**Figure 6.**
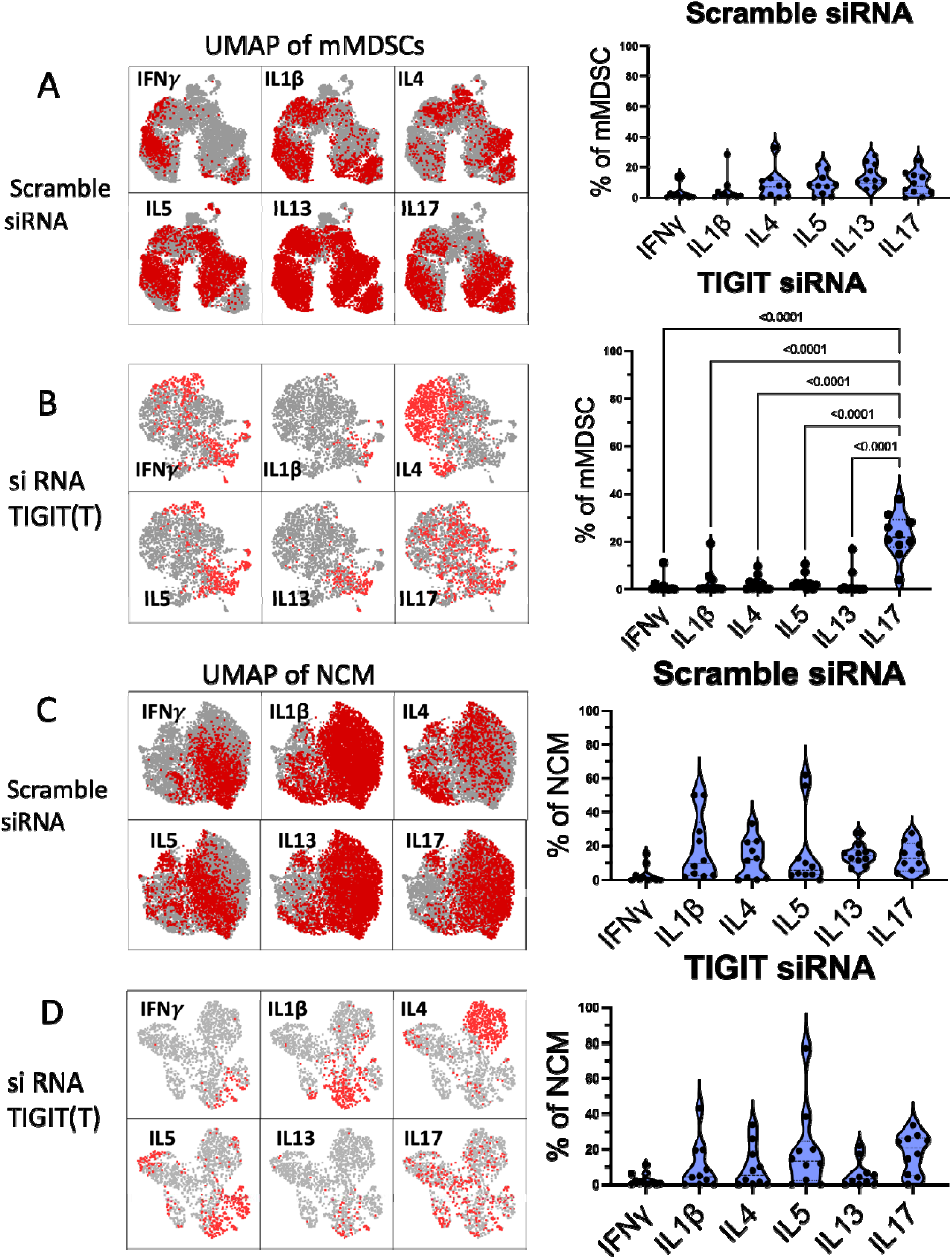
Immune modulatory milieu by immune suppressive monocytes post-TIGIT knockdown. CD11b^+^ cells isolated from healthy peripheral blood mononuclear cells (PBMCs) were knockdown by TIGIT-specific or scramble siRNA under hypoxic condition in serum free medium. These healthy monocytes were co-cultured with TEVs for 72 hours and processed for flow cytometry. (**A and B**) UMAP plots showing the cellular distribution of mMDSCs based on their expression of various cytokines (IL17, IL13, IL10, IL5, IL4, IL1β, and IFNγ) producing cells in scramble-treated (a) and TIGIT-knockdown monocytes (b). Red color represents high expression, whereas gray indicates the total cellular population of mMDSC. Violin plots comparing the percentage of mMDSCs expressing these cytokines between the scramble-treated and TIGIT-knockdown groups. (**C and D**) UMAP showing the expression of the same cytokines producing cells, (IL17, IL13, IL10, IL5, IL4, IL1β, and IFNγ) among non-classical monocytes (NCM), in scramble-treated and TIGIT-knockdown groups. Violin plots comparing the percentage of NCMs expressing various given cytokines between the scramble-treated and TIGIT-knockdown groups. Data are presented as mean□±□s.e.m. and were analyzed by one-way analysis of variance (ANOVA) with Tukey’s multiple comparisons.

### TIGIT knockdown in EV-conditioned monocytes rescues T cell proliferation in an NLRP3-dependent manner

To assess the impact of TIGIT knockdown on the immunosuppressive function of immunomodulatory monocytes, we co-cultured donor-matched T cells with EV-conditioned monocytes with or without siRNA-mediated TIGIT knockdown as described above. Our previous work has demonstrated that EV-induced immunosuppressive monocytes readily suppress T cell proliferation (*7, 10, 14*). In the present study, we found that TIGIT knockdown prior to EV conditioning rescued T cell proliferation to levels comparable to monocytes treated with scramble controls (p<0.0265, **Fig. 7A**). This also resulted in reduction of the number of IL4^+^ CD4^+^ T cells following co-culture (p<0.025, **Fig. 7D**).

**Figure 7.**
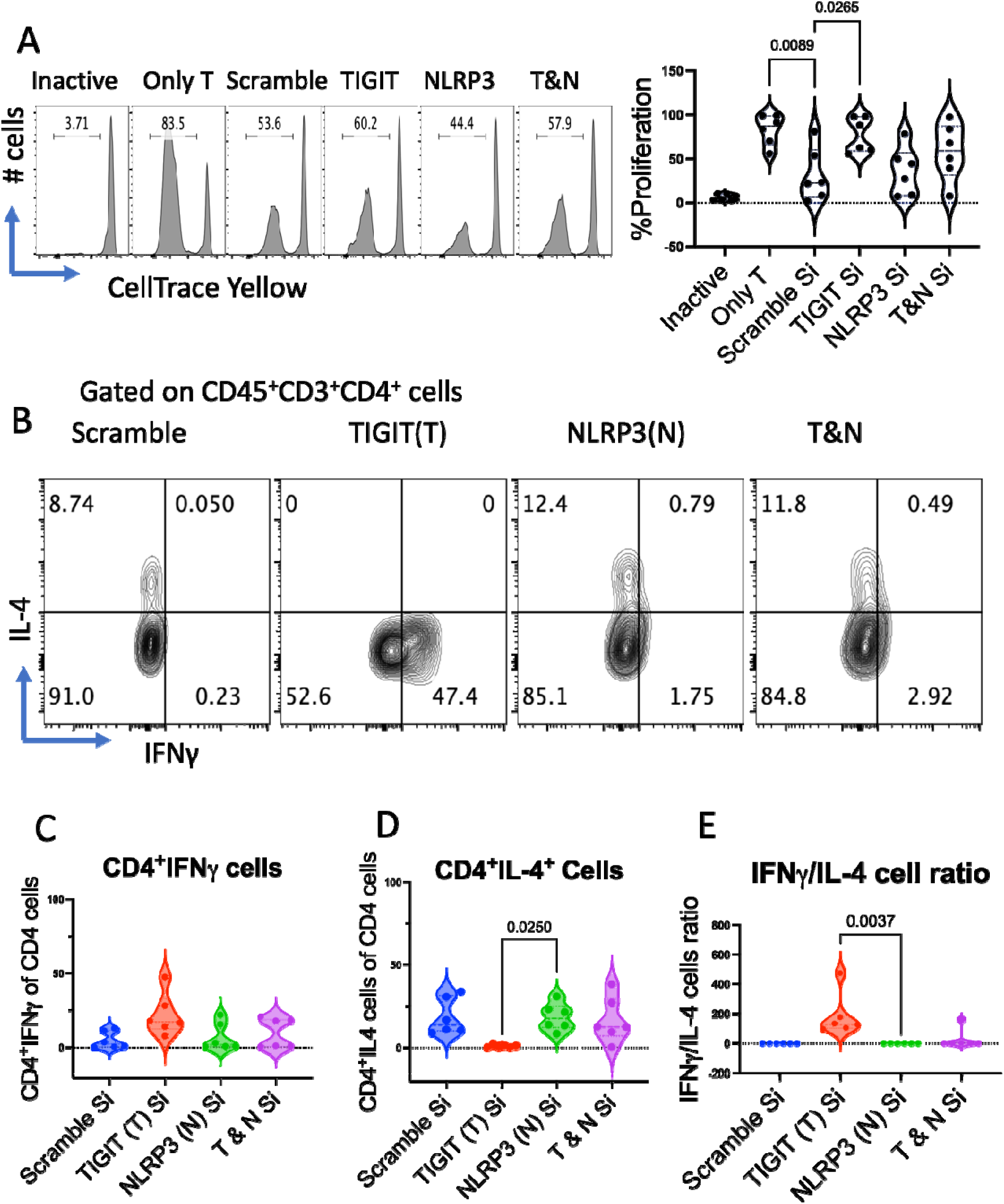
Effect of TIGIT knockdown on T Cell Proliferation in the Presence of TEV-Induced MDSCs. CD11b+ and CD3+ cells were isolated from healthy volunteer peripheral blood monocytes. CD11b cells were conditioned with TIGIT and NLRP3-specific siRNA in serum-free medium followed by priming with TEVs for under hypoxic conditions (1%) for 72 hours. Then, CD11b and T cells (T cells stained with cell trace yellow dye) were co-cultured for another 72 hours in RPMI-1640 medium with 10% serum at 37°C in 5% CO2. Finally, cells were processed for flow cytometry. (**A**) Representative flow cytometry plots and corresponding bar graphs showing CD3^+^ T cell proliferation (Cell Trace Yellow) in the absence of T cell activator, CD3/CD28 Dynbeads (inactive group), when T cells were cultured without conditioned monocytes Only T cell group), scramble treated group, TIGIT specific siRNA treated group (TIGIT), NLRP3 specific siRNA treated group (NLRP3) and both TIGIT along with NLRP3 siRNA treated group (T&N). (**B**) Representative flow cytometry plots showing IFNγ and IL-4 producing CD4+ T cells in CD3^+^ T cells and CD11b^+^ monocytes co-culture conditions, post scramble, TIGIT and/or NLRP3 knockdown. (**C**) IFNγ producing CD4+ T cells in the above-mentioned co-culture conditions in the given treatment groups. (**D**) IL-4 producing CD4+ T cells in CD3^+^ T cells and CD11b^+^ monocytes co-culture conditions, post scramble, TIGIT and/or NLRP3 knockdown. (**E**) Ratio of IFNγ producing CD4^+^ T cells and IL-4 producing CD4^+^ T cells in the given co-culture conditions. Data are presented as mean□±□s.e.m. and were analyzed by one-way analysis of variance (ANOVA) with Tukey’s multiple comparisons.

Based on our earlier results demonstrating high level of NLRP3 co-expressed with TIGIT cells, we wished to interrogate if this apparent rescue of TH1-type immune function was dependent on NLRP3 expression. By treating PBMC-derived monocytes with siRNA targeting NLRP3 and subsequently both TIGIT and NLRP3, we found that NLRP3 knockdown was sufficient to eliminate the rescue of T cell proliferation observed following TIGIT knockdown (**Fig. 7A**). NLRP3 and TIGIT double knockdown also eliminated the increase in CD4^+^ T cells observed following TIGIT knockdown (**Fig. 7B**) as well as the reduction in IL-4+ CD4+ T cells observed with TIGIT knockdown alone. Although there was not much effect on IFNγ level (**Fig. 7C**), however, the ratio of IFNγ/IL-4+ T cells increased significantly with TIGIT knockdown in monocytes alone (**Fig. 7E**), an effect that was eliminated with NLRP3 knockdown.

## Discussion

Anti-cancer immunotherapy, including immune checkpoint blockade, has revolutionized the treatment of several solid tumors in recent years (*22, 23*). Metastatic melanoma, which once carried a prognosis comparable to GBM, has seen a revolution in outcomes with immune checkpoint therapy (*3, 24*). Several clinical trials have explored these treatments in GBM, but the results have been disappointing (*5, 25, 26*). Several mechanisms for these failures have been implicated, including the mutational burden of GBM, the immune specialization of the central nervous system, but the profound immune derangement found in GBM patients is a long-described and under explored phenomena (*7, 27–29*). The mechanisms of systemic immune suppression generated by a localized disease such as GBM remains an area of active study, but induction of distal effector signals, including EVs and immunosuppressive myeloid cells, are potent avenues by which such effects could be achieved.

Work by our group and others has contributed to a growing body of evidence that myeloid cells in the GBM microenvironment are major intermediaries of tumor-mediated immune suppression, and that GBM-EVs are an effective means of inducing the formation of immunosuppressive myeloid cells (*10, 17, 30*). These myeloid cells are a highly heterogenous milieu, comprised of as many as 14 subtypes of cells as recently described (*8*), and the relative import of different immunosuppressive myeloid cell populations to the overall milieu is an open question. In our study, we focused on mMDSCs and NCMs as exemplar immunosuppressive monocyte populations, as our *in situ* tumor microenvironment data demonstrates a robust presence in the GBM microenvironment and previous work has demonstrated that these populations are modifiable by GBM-EVs (*10, 14, 30*). A central question to unlocking the complexity of the tumor-immune microenvironment is understanding the relative import and/or functional redundancy of these immunosuppressive cell populations, in order to define the critical therapeutic targets in order to effectively repolarize the tumor microenvironment to allow for anti-tumor immunity.

In the present study, we were surprised to find broad TIGIT expression across a large swath of cells in the GBM microenvironment, including immunosuppressive myeloid cells such as MDSCs and NCMs. Further, TCGA data indicated an association of TIGIT expression with reduced patient survival, indicating a potential role for TIGIT expression in disease severity. TIGIT has been classically described as a T cell exhaustion marker (*31, 32*), but some reports do describe a role for TIGIT expression in myeloid cells (*33–35*), including a study that indicated a correlation with prognosis in GBM patients (*20*). Anti-TIGIT antibodies have had some success in clinical trials as an anti-cancer immunotherapy (*36*), but have yet to show benefit in GBM treatment (*37*).

We found in our study that TIGIT expression correlated with several pro-inflammatory markers in otherwise phenotypically immunosuppressive myeloid cells, with particularly strong co-expression of TIGIT and NLRP3. NLRP3 is typically associated with the inflammasome and the production of pro-inflammatory cytokines such as IL-1B (*38–40*). The role of NLRP3 in cancer is more controversial, with many studies pointing to an immunosuppressive role, while others suggest a beneficial role (*41, 42*). The larger tumor-immune context may well provide an explanation for these mixed findings. Indeed, we found that knockdown of TIGIT resulted in a pro-inflammatory polarization of mMDSCs and NCMs, and that this effect was lost upon concomitant knockdown of NLRP3. These results are suggestive of a hierarchical arrangement of immunomodulatory signals in myeloid cells in the GBM tumor microenvironment. Inhibition of a specific immune checkpoint may be insufficient to eliminate the immunosuppressive effects of these cells given the presence of redundant signals, but prevention certain of these signals may signify a globally immunosuppressive phenotype. Prevention of the expression of certain hierarchically important signals, such as TIGIT in the present study, may be sufficient to repolarize these cells to a pro-inflammatory phenotype, which could have major implications for reversal of tumor-mediated immune suppression.

## Materials and methods

### Study design

To identify potential therapeutic targets within the glioblastoma (GBM) tumor microenvironment, we analyzed the immune phenotype of myeloid cells obtained from surgical resection specimens using ultrasonic aspirates. This direct approach allowed for the identification of immunomodulatory markers across diverse myeloid cell populations. These findings were then validated using an *in vitro* model of immunosuppressive monocyte induction, where healthy monocytes were exposed to GBM-derived extracellular vesicles (EVs) from patient-derived cell lines. TIGIT knockdown studies were then performed to interrogate its import on the immunosuppressive polarization of myeloid cells. Subsequent donor-matched T cell proliferation studies were then performed as a functional measure of immunosuppressive activity.

### Ultrasonic aspirate collection

Patients were identified with a likely diagnosis of GBM based on preoperative magnetic resonance imaging (MRI) of the brain. Written informed consent was obtained from all surgical candidates prior to surgery, in accordance with the protocol approved by the Institutional Review Board of Montefiore Medical Center (IRB# 2022-14518). Study patients (n= 14) underwent surgery with the goal of maximal safe resection. Ultrasonic aspirate was collected for research purposes based upon frozen section preliminary diagnosis of high grade glioma and included for data analysis upon confirmed diagnosis of GBM based on final histopathology and genetic testing.

Ultrasonic aspirates were immediately placed on ice for processing following collection of sample in the course of surgical resection. Fresh tumor tissue was mechanically dissociated and then enzymatically digested with TrypLE Express (Gibco, 12605010) at 37°C for 10 minutes. The resulting cell suspension was filtered through 40 µm nylon mesh, treated with red blood cell lysis buffer (BioLegend, 420301), and washed with phosphate-buffered saline (PBS). Single-cell suspensions were then prepared for flow cytometry analysis.

### Cell Culture

Patient-derived GBM cell lines BT114, BT116, and BT120, were obtained from Mayo Clinic and maintained as neurospheres in serum-free, HEPES-modified DMEM/F12 medium supplemented with EGF, FGF, and N2, as previously described.(*43*) Subsequently, these cell lines were differentiated to monolayer culture for the purposes of EV generation (dBT114, dBT116, and dBT120) in HEPES-modified EmbryoMax DMEM/F12 medium (DF-042-B, Millipore, Burlington, MA) containing 1% penicillin/streptomycin and 10% fetal calf serum. Cells were maintained in T75 flasks at 37°C with 5% CO2 in a humidified incubator, as previously described.(*10*) The cells were grown as an adherent monolayer and passaged when confluency was around 90%. Early passages within 10 or lower were used for the study.

### EV Isolation and protein estimation

For isolation of EVs, dBT114, dBT116, and dBT120 cells were transferred to serum-free neurosphere culture, DMEM/F12 medium containing EGF and FGF growth factors in 5% CO_2_ incubator. After 72hrs, cultured supernatants were collected in 50 ml centrifuge tubes and centrifuged at 1900 *g* for 10 min at room temperature. Then supernatants were transferred to 14 ml propylene Ultracentrifuge tubes, 14 × 95 mm (Cat# 331374, Beckman Coulter, Brea, CA) and centrifuged at 24000 rpm for 16 hrs at 4°C on Optima XE-90 ultracentrifuge (Beckman Coulter, Brea, CA). After centrifugation, supernatant was aspirated until ∼200μl of EV-enriched concentrated remained, consistent with prior protocols.(*10*) EVs were collected transferred to a cryovial, and protein concentration was calculated by modified Pierce Rapid Gold BCA Protein Assay Kit (A53226, Thermo Fisher, Waltham, MA) according to manufacturer’s method and the read using an Infinite 200Pro plate reader (Tecan, Mannedorf, Switzerland) using software Tecan I-Control (version 2.0). The isolated EVs were stored at −80°C until used.

### EV characterization

EVs were characterized using the ZetaView Nanotracker system (ZetaView, Particle Metrix GmbH). Freshly isolated EVs were diluted 1000-fold with filtered (40 nm syringe filter) phosphate-buffered saline (PBS) to achieve a particle concentration suitable for analysis and to remove larger particles or aggregates, and 1µL was loaded into the capillary system. The included software calculated EV size using the Stokes-Einstein equation and histograms demonstrating the distribution of EV size generated, along with movies demonstrating EV morphology.

### Peripheral blood mononuclear cells (PBMC) isolation

Human blood samples were obtained from the New York Blood Center from healthy donors (n=10). Either heparin or EDTA was used as an anticoagulant. Blood samples were delivered at room temperature (∼24°C) and processed immediately after the delivery. All subsequent steps were performed at room temperature. PBMCs were isolated from whole blood using standard density gradient centrifugation with Ficoll-Paque PLUS (17-5442-03, density 1.077 ± 0.001 g/ml, GE Healthcare, Chicago, IL) as described.(*30*) Briefly, blood was diluted 1:1 with 1X PBS and mixed gently. Ficoll-Paque™ Premium was added to the bottom of a sterile 50 ml centrifuge tube. The diluted blood was carefully layered on top of the Ficoll-Paque, maintaining a clear interface between the two layers. The tube was centrifuged at 500g for 30 minutes without brake. PBMCs were collected from the interface between the plasma and Ficoll layers using a sterile transfer pipette. The collected PBMCs were washed with PBS to remove any residual Ficoll or other contaminants and resuspended in PBS + 2% FBS. Cells were counted using the trypan blue exclusion method on Countess 3 (Invitrogen, Waltham, MA).

### Isolation of myeloid cells

Myeloid cells were positively selected from healthy donor PBMCs, using CD11b MicroBeads (130-049-601, Miltenyi Biotec, Bergisch Gladbach, Germany). Briefly, 5 µL of CD11b MicroBeads were added per 10^7^ total cells, and the mixture was incubated for 30 minutes at room temperature. Cells were then washed twice with PBS + 2% FBS and resuspended in PBS + 2% FBS. The cell suspension was applied to an LS column placed in a magnetic field of a MACS Separator (Miltenyi Biotec, Bergisch Gladbach, Germany). Unlabeled cells were collected as the flow-through fraction. The column was then removed from the magnetic field, and labeled monocytes were eluted by flushing the column with PBS + 2% FBS. The purity of the isolated myeloid cells were assessed by flow cytometry using anti-CD45 and anti-CD11b antibodies.

### Myeloid cell culture and EV treatment

Isolated CD11b^+^ cells were cultured in L-glutamine and HEPES-modified DMEM medium supplemented with 1% penicillin/streptomycin under hypoxic conditions (1% O2). Cells were seeded at a density of 1 million cells per well in a 96-well plate. Cells were treated with EVs (20 µg/well) isolated from glioblastoma cell lines (dBT114, dBT116 and dBT120) for 72hrs.

### Flow cytometry

Cultured and EV-conditioned myeloid cells were washed twice with PBS and resuspended in 50µl of PBS and processed for flow cytometric study as previously described.(*44*) Briefly, cells were stained with LIVE/DEAD Fixable Blue Dead Cell Stain (Thermo Fisher Scientific) for 30 minutes at room temperature in the dark. Following staining, cells were washed twice with FACS buffer (containing 2% human serum) and resuspended in 50µl of FACS buffer. To block non-specific binding, cells were incubated with Fc blocking solution for 20 minutes at room temperature. Subsequently, cells were washed twice with FACS buffer and resuspended in 50µl of staining buffer containing a cocktail of fluorochrome-conjugated antibodies specific for surface markers of interest. Cells were incubated for 60 minutes at 4°C in the dark. Following surface staining, cells were washed twice with FACS buffer and resuspended in 50µl of FACS buffer containing 2% human serum. Cells were fixed and permeabilized using Cytofix/Cytoperm solution (BD Biosciences, San Jose, CA) for 20 minutes at 4°C. Cells were then washed twice with permeabilization buffer and incubated with 50µl cocktail of fluorochrome-conjugated antibodies specific for intracellular markers of interest for overnight (12-14 hrs) at 4°C in the dark. After incubation, cells were washed twice with permeabilization buffer, resuspended in 200µl of FACS buffer, and filtered through a 70-micron nylon mesh to remove cell clumps. Finally, samples were acquired on a Cytek Aurora 5 laser flow cytometer and analyzed using FlowJo software.

### T cell isolation and proliferation assay

PBMCs were isolated from healthy human donors as described above. CD3+ T cells were then purified from PBMCs via magnetic cell sorting using a commercially available kit (130-050-101, Miltenyi Biotec, Bergisch Gladbach, Germany), following a similar procedure to CD11b isolation described above. Isolated CD3+ T cells were cultured at a density of 1 x 10^7 cells/ml in RPMI-1640 medium supplemented with 1% penicillin-streptomycin and 10% fetal bovine serum for 3 days (or five days in the case of siRNA knockdown experiments) at 37°C in 5% CO2. CD3+ T cells were labeled with CellTrace Yellow (Thermo Fisher) according to the manufacturer’s instructions. Autologous donor-matched labeled CD3+ T cells were co-cultured with EV-conditioned CD11b+ myeloid cells at a 1:1 ratio (1 x 10^6 cells each) in RPMI-1640 medium supplemented with 1% penicillin-streptomycin and 10% fetal bovine serum in the presence of aCD3/aCD28 DynaBeads (11131D Thermo Fisher, Waltham, MA) for 5 days at 37°C in 5% CO2. After 5 days of co-culture, cells were harvested and processed for flow cytometry analysis.

### siRNA knockdown in CD11b^+^ myeloid cells

Purified CD11b+ myeloid cells (10^6^ cells/well) were plated in a 96-well plate in 100μL of serum-free medium (Dharmacon, Lafayette, CO) containing 1% penicillin-streptomycin and transfected with Accell small interfering RNA (siRNA; Dharmacon, Lafayette, CO) at a final concentration of 1μM. Cells were incubated under normoxic conditions (37°C, 5% CO_₂_) for 48 hours. Following incubation, CD11b+ monocytes were stimulated with 20μg of tumor-derived extracellular vesicles (TEVs) and cultured for an additional 72 hours under hypoxic conditions (1% O_₂_, 5% CO_₂_, 37°C).

### Immunofluorescence staining

CD11b^+^ myeloid cells isolated from PBMCs as described above were cultured in 8-well chamber slides at a density of 1×10^6^ cells per well and allowed to adhere under standard conditions (37°C, 5% CO_₂_). After adherence, the culture medium was aspirated, and cells were gently rinsed with PBS. Fixation was performed using 4% paraformaldehyde (PFA) in PBS for 15 minutes at room temperature (RT), followed by three five minute PBS washes. Cells were permeabilized using 0.1% Triton X-100 in PBS for 10 minutes at RT and washed three times with PBS. To block non-specific binding, cells were incubated in blocking buffer (5% bovine serum albumin (BSA) in PBS) for 1 hour at RT in a humidified chamber. Cells were incubated with primary antibody diluted in blocking buffer at the recommended concentration overnight at 4°C in a humidified chamber. After incubation, cells were washed three times with PBS (5 minutes each). A fluorophore-conjugated secondary antibody was diluted in blocking buffer and applied to the cells for 1 hour at RT in the dark. Unbound antibody was removed by washing three times with PBS (5 minutes each). Cells were incubated with 1μg/mL DAPI in PBS for 5 minutes at RT and subsequently washed three times with PBS. The chamber walls were carefully removed, and a drop of mounting medium was applied to each well. Coverslips were placed gently to avoid bubble formation. Slides were allowed to set for at least 30 minutes at RT in the dark before imaging. Immunofluorescence was visualized using a fluorescence microscope equipped with appropriate filter sets.

### Western blotting

Myeloid cells, cultured with or without EVs stimulation were harvested and washed with ice cold PBS twice. Cells were then lysed in RIPA buffer supplemented with protease and phosphatase inhibitors and incubated on ice for 30 min with intermittent vortexing. Lysates were centrifuged at 12,000g for 7 min at 4°C, and the supernatant was collected. Protein concentration was determined using a Pierce Rapid Gold BCA Protein Assay Kit (A53226, Thermo Fisher, Waltham, MA) according to manufacturer’s instructions and read using an Infinite 200 Pro plate reader (Tecan, Mannedorf, Switzerland) using software Tecan I-Control (version 2.0). Equal amounts of protein (20–40 µg) were resolved on a 10–12% SDS-PAGE gel at 100V until the dye front reached the bottom. Proteins were transferred to a PVDF or nitrocellulose membrane using iBLOT3 (Invitrogen, Waltham, MA). Membranes were blocked with 5% BSA or 5% nonfat milk in Tris-buffered saline with Tween 20 (TBST) for 1 hour at room temperature. The membranes were incubated overnight at 4°C with anti-TIGIT primary antibody (PA5-121546, Invitrogen, Waltham, MA) followed by three washes with TBST (5 min each). Antibodies against HSP90 (Thermo Fisher, Waltham, MA) were used as loading control. HRP-conjugated secondary antibody was applied for 1 hour at RT, followed by three additional washes with TBST. Proteins were detected using an enhanced chemiluminescence (ECL) reagent, and signals were visualized using a chemiluminescence imaging system.

### Statistics

For statistical analysis, GraphPad Prism V.10.0 software (La Jolla, CA) was used to graph the data and perform necessary calculations. Statistical comparisons were primarily conducted using one-way analysis of variance (ANOVA) with Tukey’s multiple comparisons. Data are presented as mean ± S.E.M. and a p value threshold of <0.05 was set for determining statistical significance across all experiments. In the summary dot plots or violin plots, each dot represents an individual human value. The p values and S.E.M are indicated on these plots through error bars.

## Supplementary Materials

The PDF file includes:

Figs. S1 to S5

Tables S1 to S2

## Author contributions

M.A. and B.T.H. conceptualized, designed the experiments and wrote the manuscript. M.A., J.I. and Minori Aoki performed the experiments. C.C., P.L. and E.E. provided the patient samples. M.A. and J.I. conducted data analysis. S.M. help in preparing dot plot in figure 4. E.E., C.G. and X.Z. provided important advice. I.F.P. provided the GBM cell lines. B.T.H. acquired funding and directed the research.

## Data and materials availability

All data associated with this study are present in the paper or the Supplementary Materials.

