## Supplementary material for "TIGIT expression dictates the immunosuppressive reprogramming of myeloid cells in glioblastoma": Supplimental materials: Supplimentary materialsBIORXIV.pdf

**One Sentence Summary:** TIGIT drives immune suppression in glioblastoma via tumor extracellular vesicles and its knockdown restores T cell function through NLRP3 activation.

The PDF file includes:

Figs. S1 to S5

Tables S1 to S2

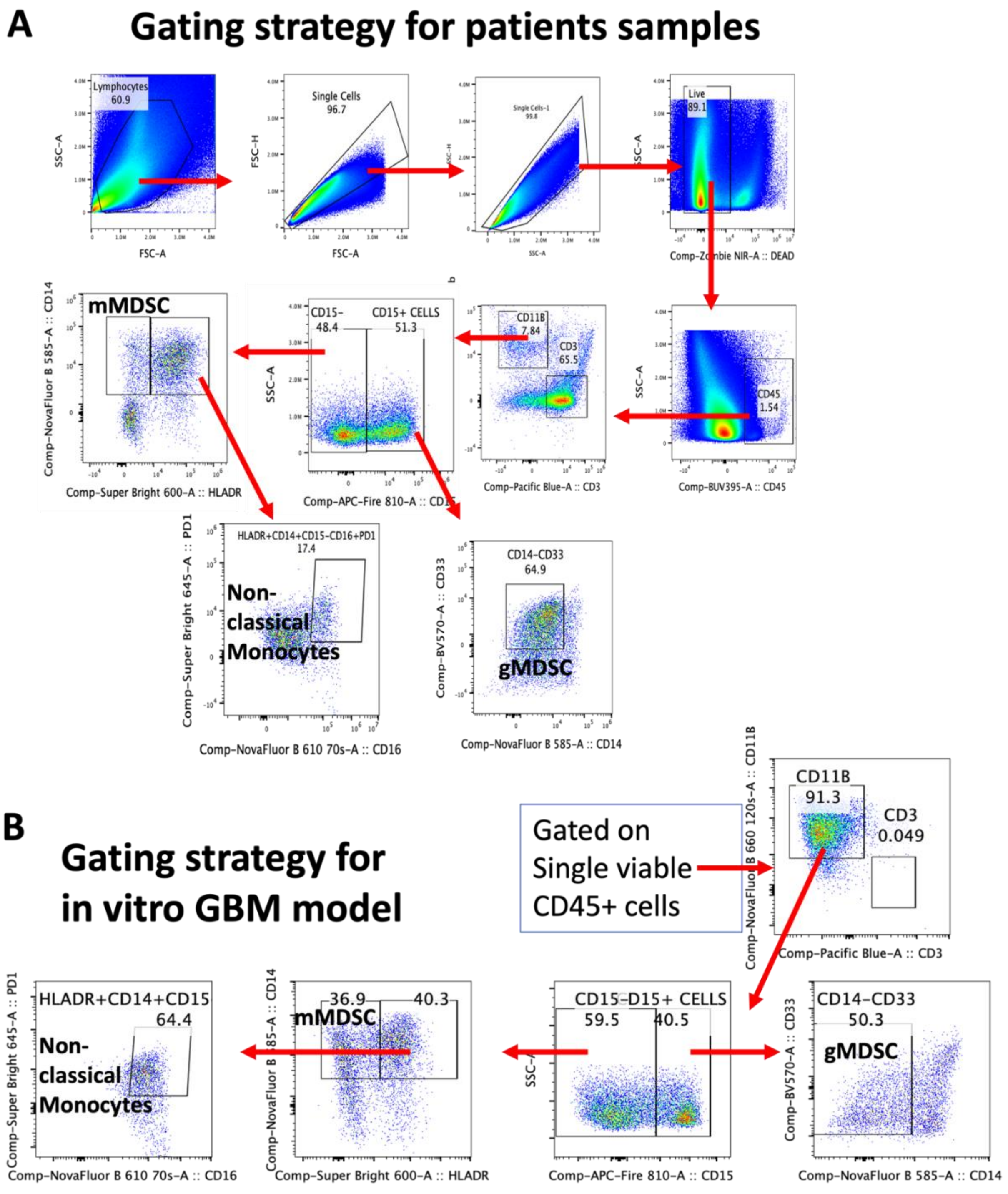

**Fig. S1 Gating strategy for the flow experiments.** Flow cytometric gating for GBM patients' ultrasonic aspirates (A). Gating strategy for in vitro GBM model of sorted peripheral blood CD11b Cells, cultured with GBM EVs (B).

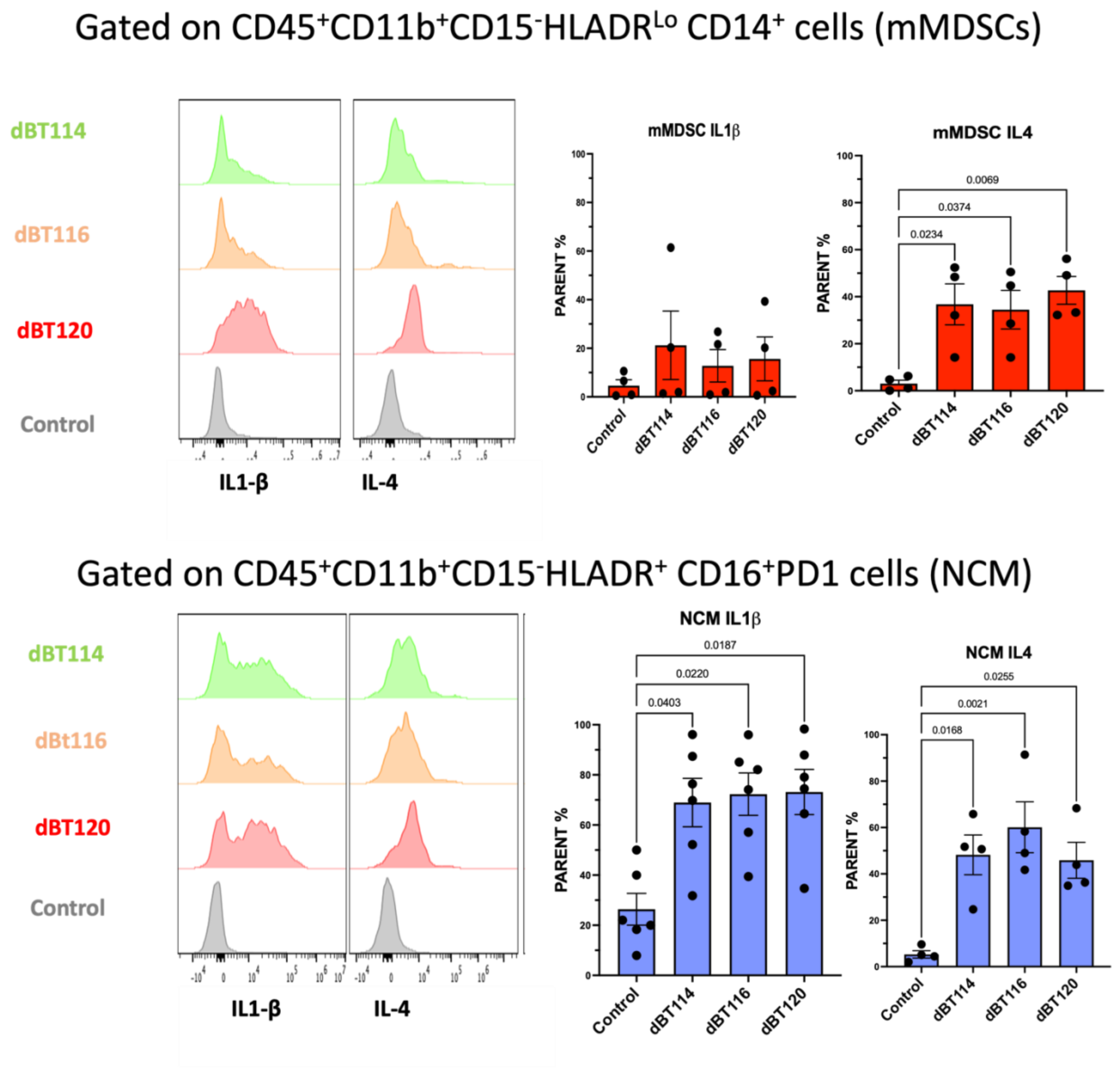

**Figure S2. Pro- and anti-inflammatory Cytokine production by suppressive monocytes post EVs treatment.** Flow cytometry analysis of IL-1β and IL-4 expressing mMDSCs (A) and NCM (B) treated with EVs obtained from dBT114, dBT116, or dBT120 GBM cells compared to no EV controls. Representative histograms illustrate cytokine expression by immunosuppressive monocytes, with quantification shown in adjacent bar graphs. Data are presented as mean ± SEM. Statistical significance was determined using a one-way ANOVA followed by Tukey's multiple comparisons test.

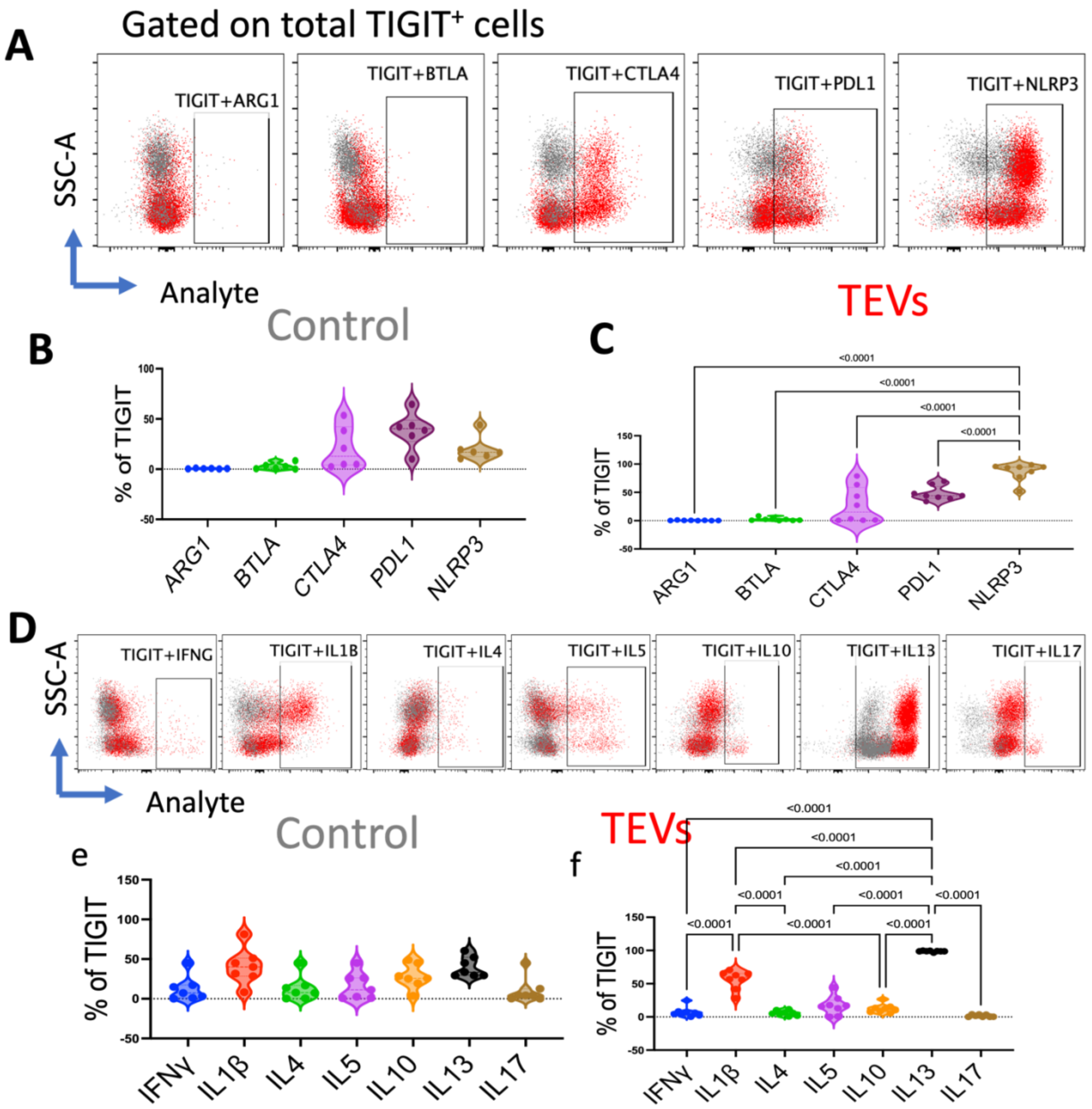

**Figure S3. TEVs induced TIGIT+ cells critically modulate the immune responses.** Flow cytometry plots showing the co-expression of TIGIT with ARG1, BTLA, CTLA4, PDL1, and NLRP3 in control and TEV-treated samples (A) and Violin plots are the summary graph for the same (B and C). Additional dot plots display the expression of various cytokines, including IFN-γ, IL-1β, IL-4, IL-5, IL-10, IL-13, and IL-17 in TIGIT+ cells (D). Violin plots summarize the distribution of cytokine expression levels (E and F). Data are presented as mean ± SEM. Statistical significance was assessed by one-way ANOVA followed by Tukey's multiple comparisons test.

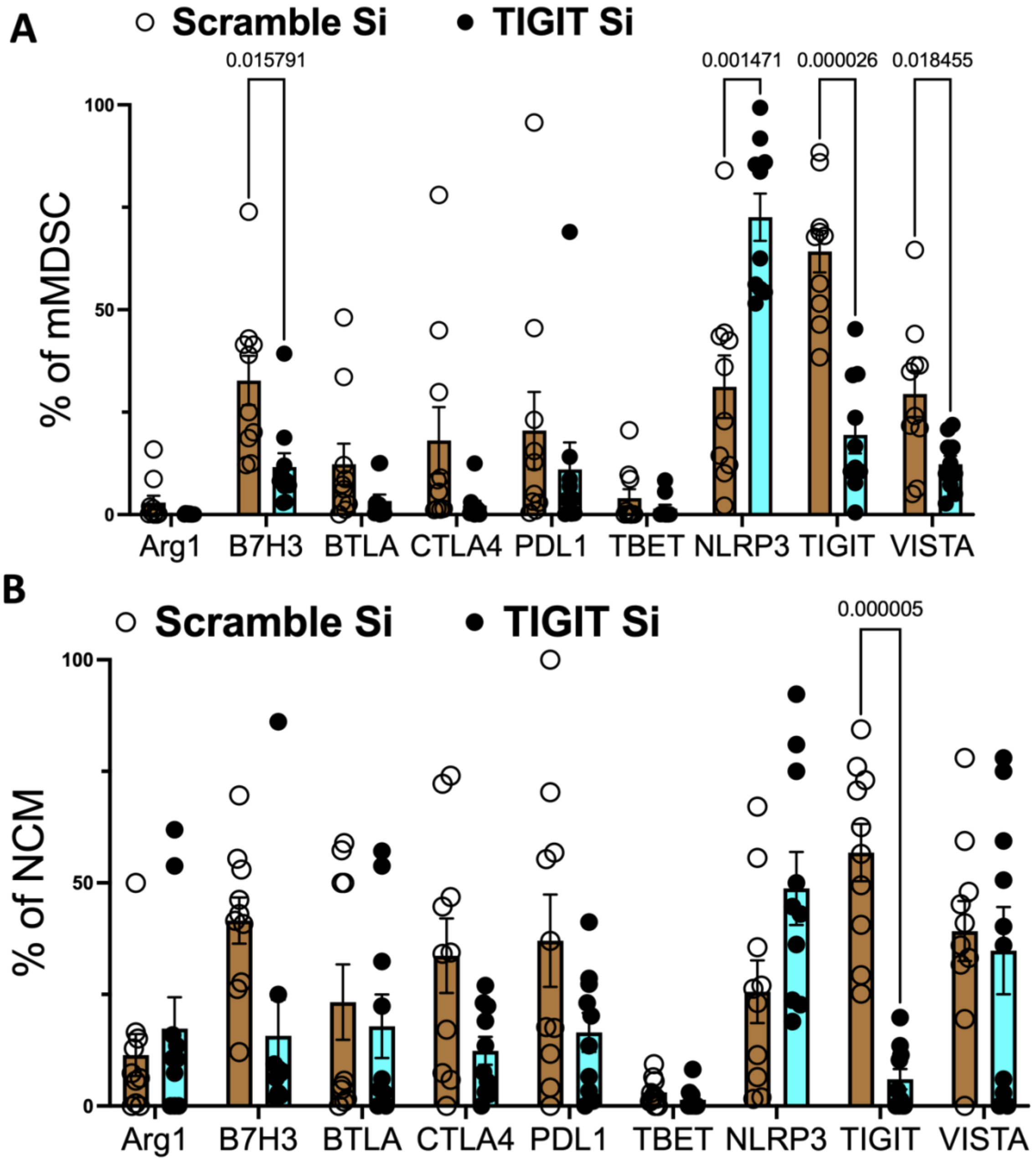

**Figure S4.** Expression of various immune modulatory molecules by mMDSCs (A) and NCM (B), post TIGIT inhibition.

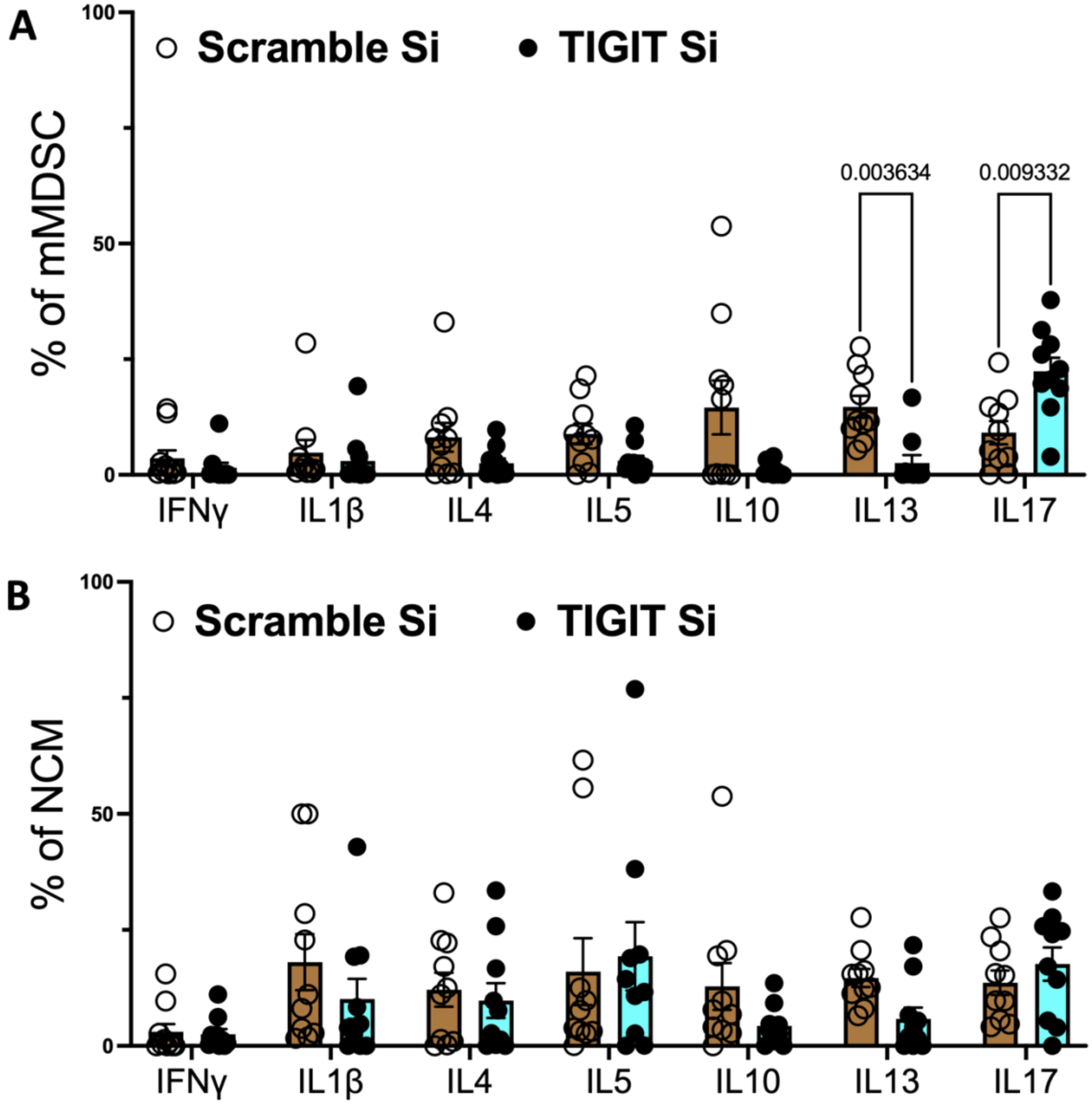

**Figure S5.** Cytokine production by mMDSCs (A) and NCM (B) post TIGIT inhibition.

#### Table S1 Patient sample Panel

| Excitation laser | Channel | Detection Filter | Marker | Clone | Fluorochrome/Dye | Peak emission |
|---|---|---|---|---|---|---|
| UV-355nm | UV1 | 373/15 | CD45 | HI30 | (BUV395) | 395 |
| | UV2 | 388/15 | | | | |
| | UV3 | 428/15 | | | | |
| | UV4 | 443/15 | BTLA | N/A | Alexa Fluor 350 | 442 |
| | UV5 | 458/15 | | | | |
| | UV6 | 473/15 | | | | |
| | UV8 | 542/28 | CD8 | RPA-T8 | 563 (BUV563) | 563 |
| | UV9 | 582/31 | | | | |
| | UV10 | 613/31 | 5-Ala | N/A | N/A | N/A |
| | UV11 | 664/27 | B7-H3 (CD276) | MIH43 | 661 (BUV661) | 661 |
| | UV12 | 692/28 | | | | |
| | UV13 | 720/29 | CTLA-4 (CD152) | 14D3 | 737 (BUV737) | 737 |
| | UV14 | 750/30 | | | | |
| | UV15 | 780/30 | | | | |
| | UV16 | 812/34 | Arginase 1 | A1exF5 | 805 (BUV805) | 805 |
| Violet-405 nm | V1 | 428/15 | IL-1 beta | AS10 | 421 (BV421) | 421 |
| | V2 | 443/15 | | | | |
| | V3 | 458/15 | CD3 | S4.1 | Pacific Blue | 450 |
| | V4 | 473/15 | IFN gamma | B27 | 480 (BV480) | 480 |
| | V5 | 508/20 | CD19 | HI19 | 510 (BV510) | 510 |
| | V6 | 525/17 | | | | |
| | V7 | 542/17 | | | | |
| | V8 | 581/19 | CD33 | WM53 | BV570 | 570 |
| | V9 | 598/20 | HLA-DR | LN3 | 600 (SB600) | 600 |
| | V10 | 615/20 | | | | |
| | V11 | 664/27 | PD1 | J105 | Super Bright 645 (SB645) | 645 |
| | V12 | 692/28 | | | | |
| | V13 | 720/29 | IL-10 | JES3-9D7 | 711 (BV711) | 711 |
| | V14 | 750/30 | PD-L1 | MIH1 | 750 (BV750) | 750 |
| | V15 | 780/30 | T-bet | 4B10 | Super Bright 780 (SB 780) | 780 |
| | V16 | 812/34 | | | | |

### Table S1 Patient sample continue

| Excitation laser | Channel | Filter | Marker | Clone | Fluorochrome/Dye | Peak emission |
|---|---|---|---|---|---|---|
| Blue-488 nm | BL1 | 508/20 | | | | |
| | B2 | 525/17 | MIF | N/A | Alexa Fluor 488 | 525 |
| | B3 | 542/17 | GFAP | N/A | Alexa Fluor 532 | 554 |
| | B4 | 581/19 | CD14 | 61D3 | NF Blue 585 | 585 |
| | B5 | 598/20 | | | | |
| | B6 | 615/20 | CD16 | SK3 | NF Blue610-70S | 610 |
| | B7 | 660/17 | CD11b | ICRF3 | NF Blue 660-120S | 660 |
| | B9 | 697/19 | CD9 | M-L13 | PerCP-Cy 5.5 | 695 |
| | B10 | 717/20 | B7x (B7-H4) | H74 | PerCP-eFluor710 | 710 |
| | B11 | 738/21 | | | | |
| | B12 | 760/23 | | | | |
| | B13 | 783/23 | VISTA | MIH65.rMAb | RB780 | 780 |
| | B14 | 812/34 | | | | |
| YellowGreen-561 nm | Y1 | 577/20 | NLPR3 | REA668 | PE | 575 |
| | Y3 | 615/20 | HHLA2 | MA57YW | PE-eFluor 610 | 610 |
| | Y4 | 660/17 | GATA3 | TWAJ | PE-Cyanine 5 | 666 |
| | Y5 | 678/18 | | | | |
| | Y6 | 697/15 | FOXp3 | PCH101 | PE-Cyanine 5.5 | 700 |
| | Y7 | 720/29 | | | | |
| | Y8 | 750/30 | CD4 | SK3 | BYG750 | 750 |
| | Y9 | 780/30 | IL-4 | 8D4-8 | PE-Cyanine 7 | 780 |
| | Y10 | 812/34 | TIGIT | VSTM3 | PE-Fire 810 | 810 |
| Red-640 nm | R1 | 660/17 | Iba1 | GT10312 | Alexa Fluor 647 | 665 |
| | R2 | 678/18 | | | | |
| | R3 | 697/19 | CD63 | H5C6 | NF Red 700 | 700 |
| | R4 | 717/20 | CD163 | GHI/61 | Alexa Fluor 700 | 720 |
| | R5 | 738/21 | Viability dye | N/A | Zombie NIR | 746 |
| | R6 | 760/23 | | | | |
| | R7 | 783/23 | IL-17A | REA1063 | APC-Vio770 | 770 |
| | R8 | 812/34 | (CD15) | W6D3 | APC/Fire 810 | 810 |

#### Table S2 In Vitro panel

| UV 349 | Marker | V 405 | Marker | B 488 | Marker | Y 561 | Marker | R 640 | Marker |
|---|---|---|---|---|---|---|---|---|---|
| UV1 373/15 | | | | | | | | | |
| UV2 388/15 | CD45 BUV 395 | | | | | | | | |
| UV3 428/15 | | V1 428/15 | IL1b BV421 | | | | | | |
| UV4 443/15 | BTLA AF350 | V2 443/15 | | | | | | | |
| UV5 458/15 | | V3 458/15 | CD3 PB | | | | | | |
| UV6 473/15 | | V4 473/15 | IFNg BV480 | | | | | | |
| | | V5 508/20 | CD19 BV510 | B1 508/20 | IL-5 Vio B515 | | | | |
| | | V6 525/17 | Cell trace Yellow | B2 525/17 | | | | | |
| UV8 542/17 | | V7 542/17 | | B3 542/17 | | | | | |
| | | | | | | YG1 577/20 | NLRP3 PE | | |
| UV9 581/31 | CD8 BUV563 | V8 581/19 | CD33 BV570 | B4 581/19 | CD14 NFB 585 | | | | |
| | | V9 598/20 | | B5 598/20 | | YG2 598/20 | RORyt RY586 | | |
| UV10 612/31 | BUV615CD11C | | | | | | | | |
| | | V10 615/20 | HLA-DR Sb600 | B6 615/20 | CD16 NFB 610-70S | YG3 615/20 | HHLA2 PE-eFluor 610 | | |
| | | | | B7 660/17 | CD11b NFB 660-120S | YG4 660/17 | | R1 660/17 | IL13 AF647 |
| UV11 664/27 | BUV665B7H3 | V11 664/27 | PD1 Sb645 | | | | | | |
| | | | | B8 678/18 | | YG5 678/18 | GATA 3 PE-Cy5 | R2 678/18 | |
| UV12 698/28 | | V12 698/28 | IL10 BV711 | B9 697/19 | CD9 PerCP-Cy5.5 | YG6 697/19 | | R3 697/19 | CD63 NFR700 |
| | | | | B10 717/20 | B7H4PERCPEF710 | | | R4 717/20 | CD163 AF700 |
| UV13 720/29 | | V13 720/29 | | | | YG7 720/29 | FOXP3 PE-Cy5.5 | | |
| | | | | B11 738/21 | | | | R5 738/21 | Zombie NIR |
| UV14 750/30 | | V14 750/30 | PD1-L1 BV750 | | | | | | |
| | | | | B12 760/23 | | YG8 750/30 | CD4 BYG750 | R6 760/23 | |
| UV15 780/30 | CTLA-4 BUV737 | V15 780/30 | Tbet SB780 | | | YG9 780/30 | IL4 PE-Cy7 | | |
| | | | | B13 783/23 | VISTARB780 | | | R7 783/23 | IL17A APC-Vio 770 |
| UV16 812/34 | ARG BUV805 | V16 812/34 | | B14 812/34 | | YG10 812/34 | TIGIT PE-Fire 810 | R8 812/34 | CD15 APC Fire 810 |
